# Vesicle-Mediated Regulation of Pollen tube Growth in *Torenia fournieri*

**DOI:** 10.1101/2024.05.19.594853

**Authors:** Xingyue Jin, Akane G. Mizukami, Satohiro Okuda, Tetsuya Higashiyama

## Abstract

In flowering plants, the growth of pollen tubes carrying sperm cells to the female gametophyte is necessary for double fertilization and seed development. The rate of growth of the pollen tube tip determines the fertility of many artificially cultivated plants and depends on the vesicle-mediated transport of cell membrane and cell wall components. However, the investigation of vesicle transport is hampered by the lack of transgenic methods for economically important plants. Here, we developed a method to transiently inhibit vesicle activity using brefeldin A (BFA) and antisense oligodeoxynucleotides (AS-ODNs) targeting key genes in the wishbone flower (*Torenia fournieri*), which has been used in studies of sexual reproduction in plants. BFA disrupted vesicle gradient homeostasis, thereby altering cell wall deposition and pollen tube morphology. The AS-ODN targeting *Torenia fournieri ANXUR* (*TfANX*), which is implicated in maintaining pollen tube growth and is used as a marker to assess inhibition by AS-ODNs, penetrated cell membranes and inhibited the expression of target genes. The treatment of pollen tubes with AS-ODN against *TfRABA4D*, which is specifically expressed in pollen and involved in regulating vesicle targeting, resulted in pollen tubes with a bulging phenotype and disrupted pectin deposition, effects similar to those of BFA. In summary, vesicle-mediated mechanisms regulate the patterning of the pollen tube cell wall in *T*. *fournieri* and our findings could facilitate the genetic manipulation of horticultural crops.

## Introduction

The delivery of sperm cells via tip-growing pollen tubes is a unique feature of seed plants, and requires precise control of tube growth during reproduction ^1^. The pollen tubes of flowering plants have high growth rates, e.g., ≤ 2.8 μm s^−^ in maize and 0.2–0.3 μm s^−^ in lily, necessitating a sophisticated vesicular trafficking system to transport secretory materials to the plasma membrane of the pollen tube tip, as well as coordination between endocytosis and exocytosis at the tip ^2–4^. Defects in tip growth are a primary cause of low fertility in artificially bred plant species ^5,6^. Therefore, investigation of the functions and mechanisms of vesicle transport in pollen tubes would provide insight into reproductive processes in angiosperms and facilitate the development of novel plant-breeding strategies.

Vesicle-mediated membrane trafficking and vesicle-regulated exocytosis and endocytosis are important for growth. In some plant taxa, such as *Arabidopsis thaliana*, maize (*Zea mays*), and tobacco (*Nicotiana tabacum*), the factors controlling vesicle trafficking have been investigated ^7–9^. Transport vesicles, including clathrin-coated, COPI, and COPII vesicles, facilitate the transport of molecules into and out of organelles and cells ^10^. Sec24, an important subunit of the COPII vesicle-coat complex in *A*. *thaliana*, recognizes and incorporates cargos into COPII vesicles ^11,12^. Disruption of its function by mutation results in reduced pollen germination and compromised transmission of the male gamete, highlighting the importance of vesicles in plant reproduction ^13,14^. SNARE proteins, which mediate vesicle fusion with the target membrane, have been characterized in *A*. *thaliana* and maize ^9,15,16^. They are classified into R-SNAREs (contributing an arginine [R] residue) and Q-SNAREs (contributing a glutamine [Q] residue). These interact to assemble a tetrameric trans-SNARE complex, thereby regulating vesicle trafficking ^17,18^. YKT61 is an R-SNARE protein in *A*. *thaliana*. Mutant ykt61*–*3 does not interact with multiple SNARE proteins and shows substantially reduced membrane association and male infertility ^19–21^. SYP121, a maize Q-SNARE protein localized to the plasma membrane, regulates K^+^ channel trafficking and cellular osmotic homeostasis ^22–24^. Rab GTPases such as RabA2 directly bind to SNARE proteins, coordinating vesicle budding and fusion in pollen tubes of *A*. *thaliana* and tobacco ^8,25–27^. Genetic interference with RabA4d, the sole member of the RabA4 subfamily expressed in pollen, leads to disturbances in wall material deposition and the formation of shorter and wider pollen tubes in *A*. *thaliana* ^27^. The functional loss of NtRab11b, a homolog of *A*. *thaliana* RabA1s, disrupts the directional growth of pollen tubes due to abnormal vesicle distribution and secretion in tobacco ^8^. Rabs regulate pollen tube morphology and growth by controlling vesicle distribution and secretion. In addition, in lily (*Lilium longiflorum*), pollen-specific formin (LlFH1) interacts with exocytic vesicles and is involved in polarized exocytosis of the pollen tube tip ^28^. These key regulators support polarized growth of pollen tubes and plant reproduction. However, the biogenesis of vesicles in pollen tubes may differ from that in other plant cells ^29^. Mutations in *A*. *thaliana VACUOLELESS1* (*VCL1*) block vacuole-related biogenesis during embryonic development, thereby modulating vesicle transport, but have minimal effects on pollen tube morphology ^30^. Furthermore, it is challenging to generate stable mutant progeny if mutant pollen tubes have severe phenotypes, such as failure to develop or germinate, abnormal tube elongation, or affected transmission of the male gametes ^31^. Hence, there is a need for alternative experimental systems that allow the investigation of key genes involved in pollen tube regulation in non-model plants.

Chemical inhibitors, which have high specificity, reversibility, and rapid efficacy, enable investigation of biochemical events in plants ^32^. For example, the fungal macrocyclic lactone brefeldin A (BFA) disrupts vesicle trafficking by inhibiting the activity of an ADP ribosylation factor G nucleotide exchange factor ^33^. On this basis, BFA has been used to investigate the secretory pathway in pollen tubes. In lily and conifer, BFA arrests pollen tube growth by disrupting vesicle trafficking and impeding the secretion of cell wall material, ultimately affecting growth polarity ^34,35^. Antisense oligodeoxynucleotides (AS-ODNs) can be used to selectively inhibit gene expression. For instance, in pear, *in vitro* pollen tube growth is hindered and reactive oxygen species (ROS) accumulation is promoted by AS-ODNs targeting periwinkle (*Catharanthus roseus*) receptor protein kinase PbrCrRLK1L13 ^36^. AS-ODNs specifically inhibit the expression of *NtGNL1* in tobacco pollen tubes, leading to abnormalities in pollen tube endocytosis ^37^. Indeed, in *A*. *thaliana* pollen tubes, phosphorothioate AS-ODNs targeting *ANX*, *callose synthase 5* (*CalS5*), and *ROP1* (a plant Rho protein), lead to growth defects, aiding the identification of genes that play an important role in reproduction ^38^. Therefore, the effects of chemicals on pollen tube growth are useful for investigating genes that affect fertility, up to and including causing infertility ^39^.

*Torenia fournieri*, an annual flowering plant, is used as a model for research on the reproduction of horticultural plants, because its naked embryo sac enables the investigation of essential fertilization events ^40,41^. Ovular attractant peptides (LUREs) and a bioactive arabinogalactan sugar chain (AMOR) are important for pollen tube guidance in *T*. *fournieri* ^42,43^. Pollen tubes of *T*. *fournieri* can be studied using microfluidics ^44^. Here, we applied BFA and AS-ODN to *T*. *fournieri* to assess the function of vesicle trafficking in pollen tube growth and its association with cell wall formation using cytochemical labelling, immunostaining, and live imaging.

BFA disrupted the distribution of vesicles in the clear zone and affected the deposition of cell wall materials, stopping tip growth. AS-ODNs entered pollen tubes in a vesicle-dependent manner and downregulated target genes, for which *ANX* was used as a marker. In addition, targeted downregulation of *TfRabA4d* by AS-ODN resulted in a shortened and bulging pollen tube, an effect similar to BFA, implicating RabA4d in vesicle trafficking. In conclusion, we used cytochemical, immunofluorescence, and molecular biological techniques to establish the AS-ODN technique in *T*. *fournieri* and characterized the functions of key genes in vesicle-trafficking during pollen tube tip growth. The method enables manipulation of the growth of the pollen tube tip, thereby paving the way for future crop enhancement.

## Results

### Inhibition by BFA disrupts vesicle distribution and pollen tube growth

To assess the function of vesicle transport in *T*. *fournieri* pollen tubes, we disrupted vesicle activity using BFA. To germinate pollen we used pollen germination medium (PGM) containing 1, 2, 3, or 4 μg/ml BFA, based on previous reports (0.1–20 μg/ml in tobacco ^35,45^, 0.1–5 μg/ml in *A*. *thaliana* ^34,46,47^, 1 μg/ml in lily ^34^, and 8 μg/ml in a conifer^35^). BFA at 4 μg/ml was sufficient to inhibit vesicle trafficking during pollen germination of *T*. *fournieri* (Fig. S1). Therefore, pollen tubes germinated in PGM without BFA for 1 h were treated with 4 μg/ml BFA for 2 h (PGM 1 h and BFA 2 h) to investigate vesicle trafficking in pollen tubes (Fig. S1).

Pollen tubes treated with BFA showed slowed growth with morphological abnormalities (Fig. 1A-D). Although most pollen tubes germinated in PGM for 1 h, those transferred to medium containing 4 μg/ml BFA showed a pollen tube germination rate of 78.1 ± 18.6%, which is lower than the 91.7 ± 3.9% in PGM for a further 2 h (Fig. 1A, B). Therefore, late germination was inhibited after 1 h of cultivation. In addition, the tips of BFA-treated pollen tubes showed a shorter and bulging phenotype (Fig. 1C, D).

**Fig. 1.**
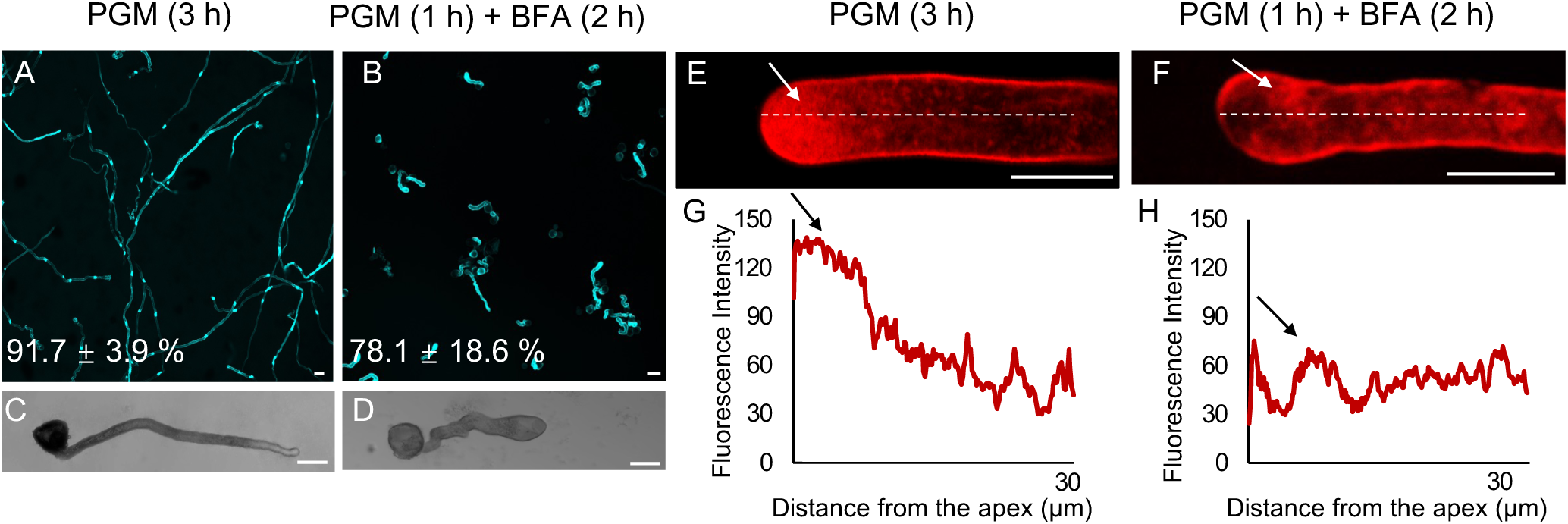
Effect of BFA on pollen germination, pollen tube morphology, and vesicle distribution in *T. fournieri*. **A** Pollen tubes cultured in standard PGM for 3 h showed a germination rate of 91.7 ± 3.9% and a normal shape. **B** Pollen tubes cultured in PGM for 1 h followed by PGM containing 4 μg/ml BFA for 2 h showed a germination rate of 78.1 ± 18.6% and bulging. **C** Normal morphology of pollen tubes cultured in PGM (3 h). **D** Typical morphology of pollen tubes cultured in PGM (1 h) and BFA (2 h). **E** Vesicle distribution in a pollen tube stained with FM4-64 in PGM (3 h). **F** Uneven vesicle distribution in a pollen tube stained with FM4-64 in PGM (1 h) and BFA (2 h). **G** Relative fluorescence intensity on the white dashed line in (**E**). **H** Relative fluorescence intensity on the white dashed line in (**F**). Arrows indicate accumulations of secretory vesicles in the apical zone (**E**, **F**); the corresponding fluorescence intensities are shown in (**G**, **H**). Accumulated vesicles in **F** (arrow) in the BFA compartment. Scale bars: 10 μm.

To determine whether the reduced growth of pollen tubes was caused by inhibition of vesicle trafficking, we used the amphiphilic dye FM4–64 to label vesicles in living pollen tubes of *T*. *fournieri*. FM4–64 consistently labeled the cell membrane and vesicles of normally growing *T*. *fournieri* pollen tubes, with intense staining in the apical region. Staining was distributed in the extreme apical region (2–10 μm); outside this region, the stain was prevalent and significantly attenuated (Fig. 1E, G). BFA-treated pollen tubes had a wider distribution of signal but intense apical staining was absent and the signal intensity was 50% of untreated pollen tubes. The signals aggregated at the subapical region of BFA-treated pollen tubes, a location termed a *BFA compartment* (Fig. 1F, H). Live-cell imaging showed a correlation between vesicle apical accumulation and pollen tube tip growth (Movies S1,2). Normally growing pollen tubes had a growth rate of 0.16 μm s^−^ and showed vesicle accumulation in the apical region (Movie S1). After treatment with PGM (1 h) and BFA (2 h), the apical region of the pollen tube showed negligible vesicle accumulation, swelling at the tip, and significantly inhibited growth, which hampered assessment of the growth rate (Movie S2).

### Disruption of vesicle distribution influences the composition of the pollen tube cell wall

The chemical composition of the cell wall supports the rapid growth of the pollen tube tip. Typically, the cell wall comprises an inner layer rich in callose (β-1,3 glucan) and an outer fibrillar layer, which mainly consists of methyl-esterified pectin. During the maturation of the cell wall, pectin undergoes demethylation, enabling the binding of calcium ions to enhance the mechanical strength of the cell wall ^48,49^.

To determine the mechanism underlying the swelling morphology in BFA-treated pollen tubes, we performed cytochemical and immunofluorescence labeling to investigate the structures of the cell membrane and cell wall (Fig. S2). Staining with the membrane-specific dye CellMask™ showed no differences in the cell membranes of control and BFA-treated pollen tubes (Fig. S2A). However, aniline blue, which stains callose, showed marked aggregation of callose in pollen tubes cultivated in PGM (1 h) and BFA (2 h) compared to untreated pollen tubes (Fig. S2A). Most pollen burst after treatment with BFA for 3 h, as indicated by the absence of tubular structures in the cell membrane and callose (Fig. S2C).

The pectin gradient in the pollen tube apex is important for tip growth; highly esterified pectin is concentrated at the extreme apex and decreased towards the base. Pectin methyl-esterase (PME) sustains this gradient by de-methoxylating pectin, enabling calcium-mediated crosslinking and enhancing cell wall rigidity ^50^. The softer apical wall facilitates rapid growth episodes, which are closely associated with pectin de-esterification. We performed immunostaining using JIM5, JIM7, and LM16 antibodies to investigate the distributions of unesterified, methyl esterified, and both pectins, respectively ^51^ (Fig. 2). In the normal pollen tube, JIM5 signal was present in the shank zone, and was negligible in the apical region. JIM7 fluorescence indicated that methyl-esterified pectin was primarily located in the apical region, as confirmed by Ruthenium red staining. This suggests a gradient of methylated and demethylated pectin along the pollen tube. LM16 staining was distributed uniformly throughout the pollen tube. The distribution of cell wall components after cultivation in PGM (1 h) and BFA (2 h) exhibited considerable aberrations. JIM5 and JIM7 displayed similar distributions, with the JIM5 staining intensity being stronger at the pollen tube tip, indicating disruption of the pectin gradient. The LM16 signal (marking basic pectin precursors) accumulated at the swelling tip.

**Fig. 2.**
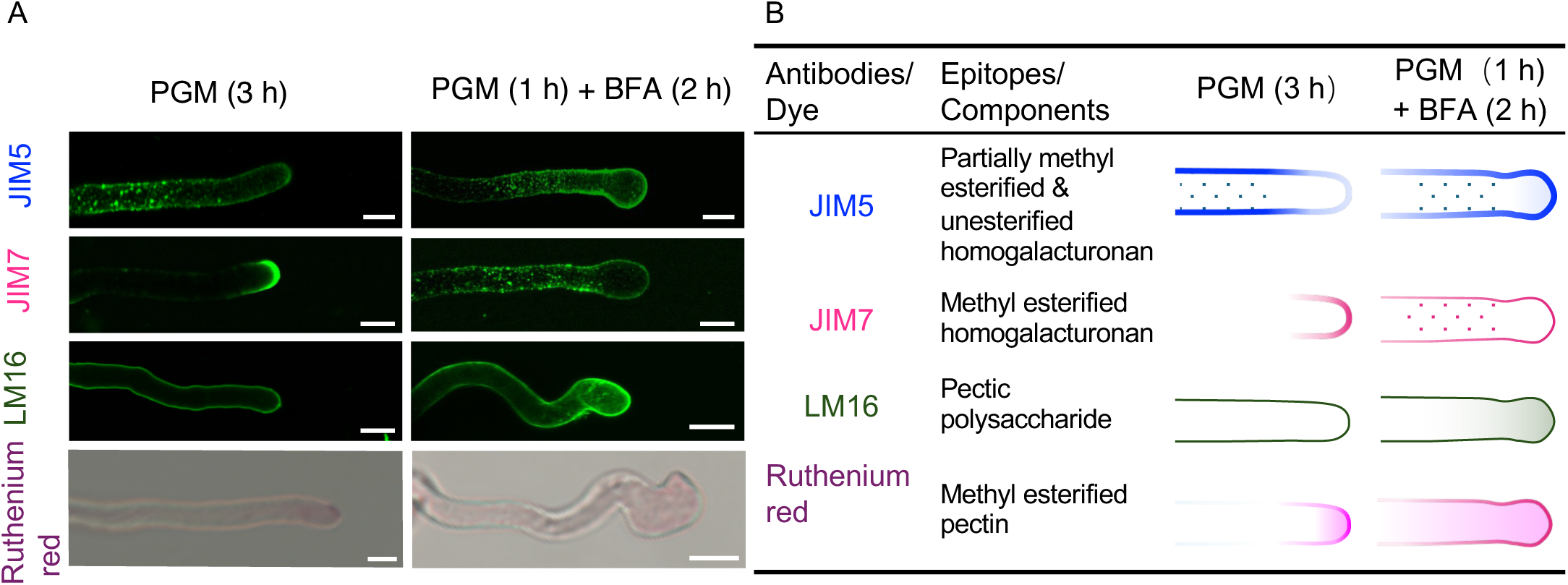
Pectin labeling in pollen tubes in the presence and absence of BFA. **A** Distribution of pectin components in pollen tubes cultured on PGM (3 h) or PGM (1 h) and BFA (2 h), labeled using JIM5, JIM7, and LM16 monoclonal antibodies, and stained with ruthenium red. **B** Summary of pectin component distributions. Scale bars: 10 μm.

### ODN enters the pollen tube from the tip in a vesicle-dependent manner

AS-ODNs have been used to modulate gene expression in pollen tubes in tobacco and *A*. *thaliana*. We first ensured that AS-ODNs could pass through the pollen tube cell wall and plasma membrane. *TfANX*-targeting ODNs labeled with AlexaFluor488 were used to stain pollen tubes germinated on PGM. After ∼1 h, Alexa 488 signal accumulated at the pollen tube tip, and small punctate Alexa 488 signals were evident in the cytoplasm (Fig. S3). After 3 h, the signal was widely distributed throughout the pollen tube, indicating penetration of ODNs (Fig. 3A, C).

**Fig. 3.**
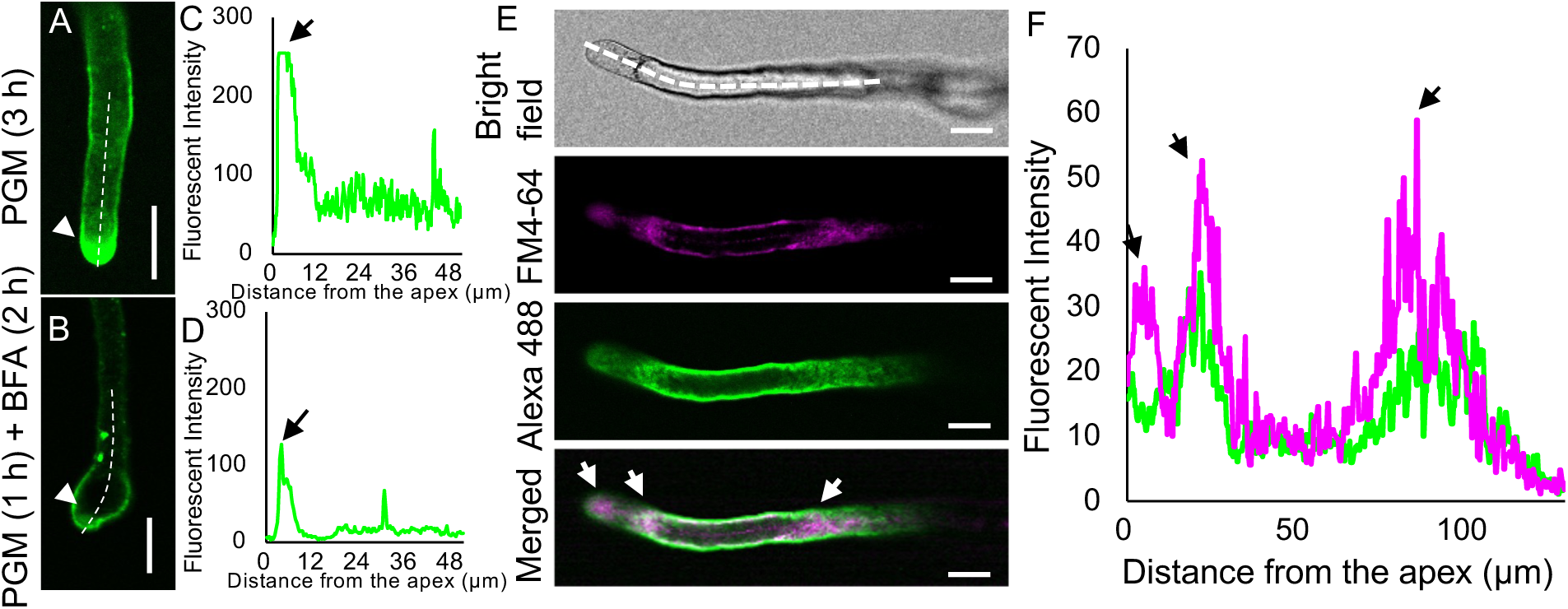
Uptake of AS-ODNs into pollen tubes. **A** Alexa488-labeled ODN entering the tip of a pollen tube cultured on PGM (3 h). **B** Alexa488-labeled ODN entering the tip of a pollen tube cultured on PGM (1 h) and BFA (2 h). **C-D** Fluorescence intensities of **A** and **B**; arrowheads, clear zones. **E** Pulse-chasing labeling with Alexa 488-labeled ODNs and FM4-64 in the same pollen tube. **F** Fluorescence intensities of **E**; arrows, clear zones and the start and end points of the vacuole. n = 20; scale bars: 10 μm.

ODNs enter pollen tubes in a vesicle-dependent manner ^37^. To determine if this was the case in *T*. *fournieri*, we visualized the uptake of Alexa 488-labeled ODNs in pollen tubes cultivated in PGM (1 h) and BFA (2 h). The Alexa488 signal notably decreased in pollen tubes cultivated in PGM (1 h) and BFA (2 h), suggesting that the inhibition of vesicle activity by BFA affected the absorption of ODNs (Fig. 3B, D).

ODN uptake first at the tip and then throughout the pollen tube is similar, but slower, than FM4–64 uptake. We investigated whether ODNs entered *T*. *fournieri* pollen tubes by vesicle-mediated endocytosis. The Alexa 488-labeled ODN and FM4–64 (to visualize endocytosis) signals overlapped at the tip (Fig. 3E) but gradually decreased in intensity and the colocalization was disrupted in the shank. The FM4–64 signal was significantly stronger than Alexa 488-labeled ODN in the large vacuolar region (Fig. 3F). The Alexa 488-labeled ODN signal did not completely overlap with that of FM4–64, which implies that some ODN escapes from endosomes into the cytoplasm.

### Inhibition of *TfANX* expression by AS-ODN alters pollen tube growth *in vitro*

To assess the effect of AS-ODN on target genes in *T*. *fournieri* pollen tubes, we designed an AS-ODN and its sense control (S-ODN) targeting *TfANX* (Figs. 4, S4) based on the conserved region shared by ANX1 and ANX2, which are implicated in the maintenance of pollen tube growth (Fig. S4). Addition of ODNs to PGM affected the pollen germination rate in a concentration-dependent manner (Fig. S5). At 1 μM, neither AS-ODN nor S-ODN significantly affected the pollen germination rate or pollen tube morphology. There were slight differences with application of 5 μM ODNs but ODNs at 10 μM were optimal. Therefore, we used 10 μM ODNs in subsequent experiments, which was lower than that used for *A*. *thaliana* (20 μM).^38^ Treatment of pollen with 10 μM ODNs significantly reduced *TfANX* expression in pollen tubes (Fig. 4B), indicating that AS-ODN is capable of modulating gene expression in *T*. *fournieri* pollen tubes *in vitro*. Furthermore, AS-ODN caused shorting (30.8%), branching (13.4%), and bursting (15.4%) of pollen tubes, which resembled those of *A*. *thaliana anx1/2* mutants ^52^. This method can be used to validate gene function, because generating knockdown and knockout mutants is problematic.

**Fig. 4.**
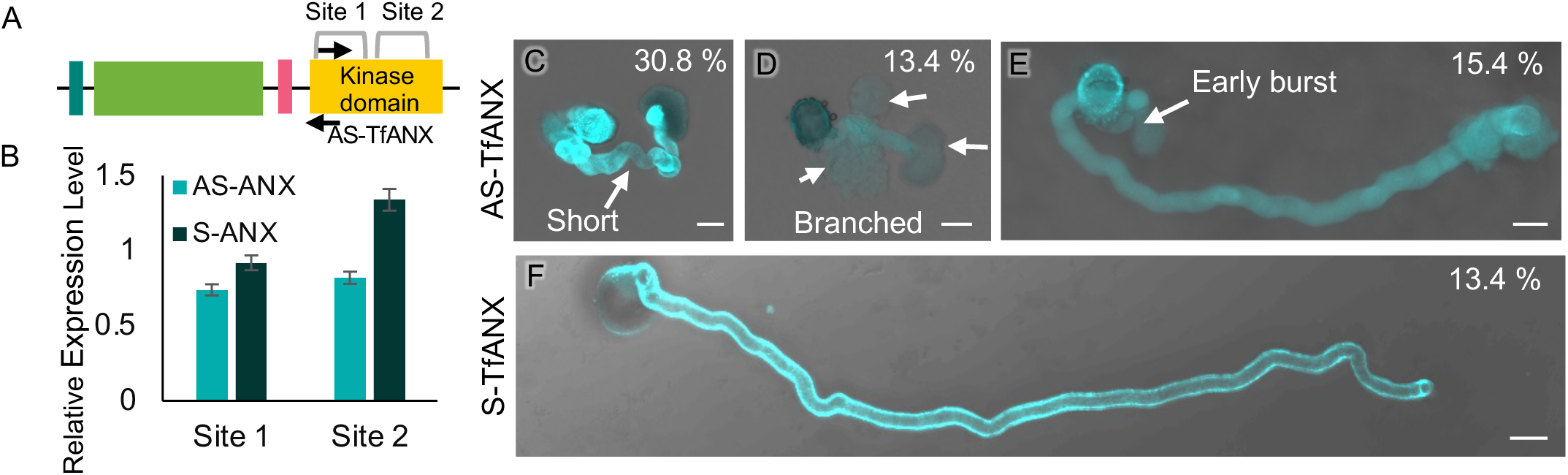
Inhibition of pollen tube growth by AS-ODNs to *TfANX in vitro*. **A** Structure of the TfANX domain and the sites of qPCR primers. Arrows show the AS-and S-ODN sites. **B** Relative expression level of *TfANX* in AS-ANX- and S-ANX-treated pollen tubes. **C-F** Treatment with AS-TfANX led to short (**C**), branched (**D**), and early burst (**E**) pollen tubes compared to the S-TfANX control (**F**). Arrows indicate phenotypes. Data are from > 300 pollen tubes. Scale bars: 10 μm.

### Modulation of *TfRABA4D* expression by AS-ODN alters vesicle dynamics and pollen tube morphology in *T*. *fournieri*

The RabA family of Rab GTPases regulates vesicle formation, motility, and tethering, particularly in membrane trafficking linked to the trans-Golgi network. The absence of RABA4D disrupts polar growth and alters cell wall patterning in *A*. *thaliana* ^27^. We investigated the role of TfRABA4D in vesicle function and pollen tube growth in *T*. *fournieri* using AS-ODNs.

TfB078315 was selected as TfRABA4D, the homologs of RABA4D, because of TfB078315 is closest to RABA4D on the evolutionary tree and its high expression in pollen. (Fig. S6).

To investigate the function of TfRABA4D, we analyzed germinated *T*. *fournieri* pollen tubes cultivated for 3 h in AS-/S-TfRABA4D medium. AS-TfRABA4D-treated tubes were shortened and had bulges at the tips, thus their morphology was different and their *RABA4D* expression was reduced compared to S-TfRABA4D-treated pollen tubes (Fig. 5B-E, J). This phenotype resembled pollen tubes treated with BFA, indicating that TfRABA4D modulates vesicle trafficking. The uptake of FM4–64 was significantly decreased in AS-TfRABA4D-treated tubes, confirming that *TfRABA4D* modulates vesicle dynamics in *T*. *fournieri* (Fig. 5F, G).

**Fig. 5.**
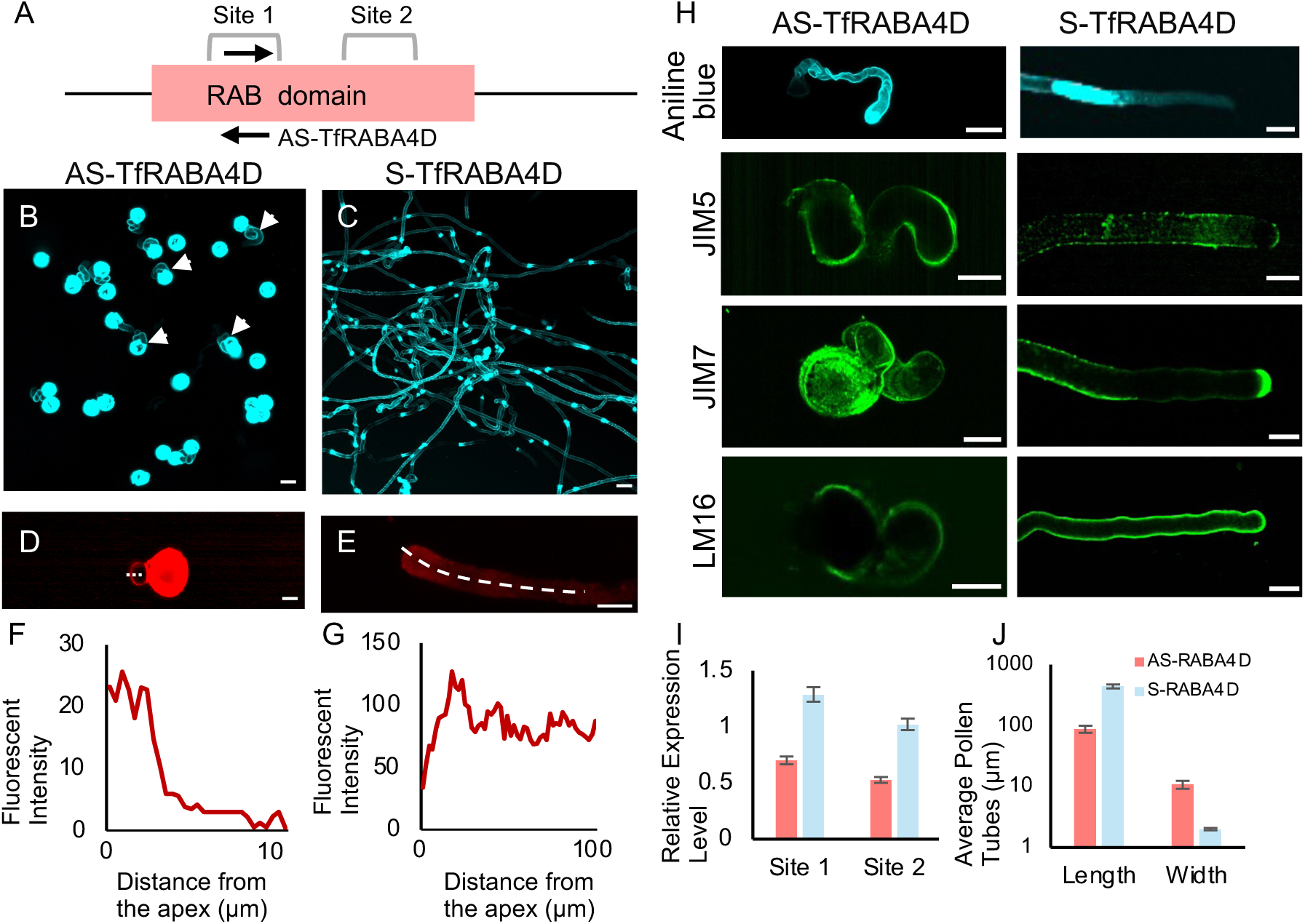
Effects of AS-ODNs to *TfRABA4D* on pollen tube morphology and cell wall patterning. **A** Domain structure of TfRABA4D; arrow, ODN site; brackets, sites of qRT-PCR primers. **B, C** Images of pollen grains germinated in 10 µM AS-TfRABA4D (**B**), and 10 µM S-TfRABA4D medium (**C**). Arrowheads indicate abnormally enlarged pollen tubes. **D, E** Image of vesicle distribution by FM4-64 staining in pollen tubes germinated after cultivation for 3 h in 10 µM AS-TfRABA4D (**D**), and 10 µM S-TfRABA4D medium (**E**). **F, G** Fluorescence intensities of **D** and **E**. **H** Callose and pectin distribution in pollen tubes treated with AS- or S-TfRABA4D and stained with aniline blue or immunostained using JIM5, JIM7, and LM16 monoclonal antibodies. **I** Relative expression levels of TfRABA4D in AS- and S-TfRABA4D-treated pollen tubes. **J** Average length and width of AS- and S-TfRABA4D-treated pollen tubes. Data are from > 300 pollen tubes. Scale bars: 10 μm.

The downregulation of *TfRABA4D* did not influence the germination rate, indicating dysregulated deposition of pollen tube cell wall components (Fig. 5H). Aniline-blue staining revealed a uniform callose distribution throughout AS-TfRABA4D-treated pollen tubes, compared to reduced callose deposition at the tip and spots of accumulation in sense-treated pollen tubes. In AS-TfRABA4D-treated pollen tubes, the distribution of unesterified homogalacturonan (JIM5 signal) was less intense. Conversely, methyl-esterified homogalacturonan (JIM7 signal), which typically accumulates at the tip in S-TfRABA4D-treated pollen tubes, was dispersed throughout the pollen tube. The distribution of pectic polysaccharides (LM16 signal) was more diffuse compared to S-TfRABA4D-treated pollen tubes. These findings implicate *TfRABA4D* in the vesicle trafficking upon which pollen tube patterning depends.

## Discussion

Chemical inhibitors have been used in molecular mechanistic investigations of pollen tubes. BFA, latrunculin B, cytochalasin D, and colchicine have provided insight into the functions of vesicle trafficking, actin polymerization, and microtubule polymerization in pollen tube growth ^32^. The novel compounds disruptol-A and disruptol-B, which may target lipid kinases, indirectly affect the tip-growth machinery in *A*. *thaliana* ^53^. In this study, we applied BFA to germinated *T*. *fournieri* pollen tubes and conducted biochemical and immunofluorescence analyses to ascertain the effect of BFA-induced inhibition of vesicle movement on pollen tube endocytosis and exocytosis. However, chemical inhibitors are toxic to pollen tubes and have multiple physiological effects, hampering the investigation of pollen tube behavior.

Applying AS-ODNs to complementary mRNA sequences has been used to suppress gene expression and investigate gene functions ^38^. However, the negative charge of ODNs hampers their passage across the plant cell wall. Therefore, cationic lipids have been used to transport ODNs ^54^. Sucrose and active transport via sugar translocators can significantly enhance AS-ODN uptake ^37^. We initially used PGM containing 5% sucrose and Alexa488-labeled AS-ODNs. An Alexa488 signal was observed in pollen tubes, indicating ODN uptake during growth. By contrast, BFA, which inhibits vesicle activity, reduced ODN uptake by pollen tubes. The overlap between Alexa488-labeled ODNs and FM4–64 at the pollen tube tip suggested that specialized membrane trafficking at the apex facilitates AS-ODN uptake. The mechanism of ODN uptake warrants further investigation, because BFA inhibits exocytosis rather than endocytosis. It is plausible that ODN escapes from endosomes to the cytosol to inhibit gene expression (Fig. 3), but the mechanism is unknown. Our method enables analysis of gene function using pollen tubes as a model system, as well as analysis of how ODN functions.

Vesicles vary in size and are distributed differently along the growing pollen tube. For example, smaller vesicles are involved in membrane recycling and larger vesicles carry bulky cell wall materials and proteins ^55^. Several regulators of vesicular trafficking have been identified in mutagenesis studies ^7,22,27^. In pollen tubes, the specialized haploid cell responsible for tip growth relies on dynamic vesicular trafficking. However, because of their functional importance, mutations in these regulators can lead to lethality, hampering in-depth mechanistic analysis ^56^.

Chemical inhibitors enable disruption of vesicular trafficking in pollen tubes grown *in*_J*vitro* without time-consuming manipulations. In *T*. *fournieri*, the pollen tube growth rate was decreased by BFA. Indeed, pollen germinated in AS-ODN medium showed abnormal growth characterized by bulging pollen tubes. Our method enables the characterization of vesicular trafficking during *T*. *fournieri* pollen tube growth, and we plan to apply it to evaluate the cellular functions of plant RABs and how the processes they mediate affect pollen growth and fertility in *T*. *fournieri*.

Rab GTPases, such as Rab11b and RabA4d, localized to the tip of pollen tubes, mediate the targeting of transport vesicles to the apical clear zone by controlling the budding and fusion of vesicles with the plasma membrane through an interaction with membrane-trafficking effector proteins ^26^. AS-ODNs targeting *TfRABA4D* caused abnormal phenotypes in *T*. *fournieri* pollen tubes, including bulging. Similar phenotypes were observed in BFA-treated pollen tubes and an *A*. *thaliana raba4d* mutant. This suggests that the phenotypes of AS-ODN-treated pollen tubes were caused by downregulation of *TfRABA4D* ^27^. Vesicle trafficking is important for the transport of pectin and other cell wall materials in pollen tubes. Pulse-chase labeling of vesicle trafficking in the pollen tubes showed that downregulation of *TfRABA4D* influenced vesicle distribution and cell wall deposition, leading to abnormal pollen tube growth and morphology. The gradient distribution of pectin methylation in pollen tubes, which is important for growth and guidance, is established by the coordinated actions of pectin methylesterase (PME) and pectin methylesterase inhibitors (PMEIs) ^57,58^. PME, which demethylates pectin, is localized to the tips of pollen tubes. The disruption of vesicle trafficking by BFA or downregulation of *TfRABA4D*, leads to loss of this gradient of pectin methylation. This suggests that vesicle trafficking is critical for the deposition of pectin and other cell wall materials in pollen tubes and that its disruption leads to abnormal growth and morphology. However, it is unknown whether the primary effect of vesicle-trafficking disruption is on the transport of cell wall materials or the distributions of PME and PMEIs. Further research should aim to identify the mechanisms by which vesicle trafficking regulates pectin methylation and pollen tube growth.

In conclusion, BFA and AS-ODN targeting *TfRABA4D* modulated vesicle activity in *T*. *fournieri* pollen tubes. This affected the morphology of pollen tubes, leading to a pectin methylation gradient in the cell wall. The findings establish a molecular basis for vesicle-trafficking dynamics. Our method can be used in studies of pollen tubes of non-model plant species.

## Materials and Methods

### Plant materials and growth conditions

*T*. *fournieri* plants were grown in soil at 25°C under long-day conditions (16 h light/8 h darkness) ^41^. Flower buds ∼2 days after anthesis were selected for pollen tube cultivation. Pollen was germinated and cultured in modified Nitsch’s medium (PGM) ^59^. The germination rate was altered in PGM containing BFA (1, 2, 3, and 4 μg/ml). Therefore, after incubation for 1 h in PGM, pollen tubes grown *in vitro* were incubated with BFA (4 μg/ml) for 2 h (BFA-treated group).

### Genome-wide identification and gene expression analysis

Seventeen *A*. *thaliana* CrRLK1L protein sequences were obtained from the TAIR website and 23 CrRLK1L proteins were identified in *T*. *fournieri* (TfCrRLK1L) based on the presence of malectin (pfam: TfB080671) and kinase (pfam: PF07714) domains using the Pfam Protein Family database (http://pfam.xfam.org/). RAB genes were identified in the same way based on the *A*. *thaliana* sequence and the RAS domain (pfam: PF00071). The expression profiles of *TfCrRLK1L* and *TfRAB* genes in different tissues and developmental stages were obtained from public transcriptome databases ^60^. We used TBtools to visualize the structures of candidate genes, protein domains, and expression profiles ^61^. The cDNA sequences of the genes used in this study were listed in Table S2.

### ODN selection and delivery to pollen tubes

ODNs were designed to target sequences predicted using the Sfold tool (https://sfold.wadsworth.org/cgi-bin/soligo.pl), based on the principles of nucleic acid thermostability ^38^. We selected multiple 21 bp antisense fragments from the mRNA of the targeted gene, which were capped with phosphorothioate at the 5’- and 3’-ends (Table S1). These sequences had high GC contents and were synthesized by Custom Primer Invitrogen™ (Thermo, Japan).

ODNs were dissolved in PGM to the desired concentrations. Pollen grains were placed on PGM and incubated for 3–6 h at 28°C in a humidity chamber. Next, germinated pollen on PGM was stained with 0.1% aniline blue. Three independent experiments were conducted on different days to evaluate the effects of the ODNs.

### Extraction of RNA from pollen tubes and qRT-PCR

RNA was extracted using the RNeasy Plant Mini Kit (Qiagen, USA). Pollen tubes were isolated by centrifugation at 3000 rpm. The supernatant was decanted and TRIzol was added following the manufacturer’s guidelines. RNA samples were normalized to uniform concentrations using the BioSpec-nano spectrophotometer (Shimadzu, Japan). RNA was reverse-transcribed using the SuperScript II Reverse Transcriptase Kit (Invitrogen, USA), following the manufacturer’s protocols. The primers listed in Table S1 were used for qRT-PCR, with GADPH as the reference gene for the 2^−ΔΔCt^ technique ^42^.

### Cytochemical labeling and immunostaining of pollen tubes

To stain vesicles, pollen tubes were incubated with 2 μM FM4–64 in PGM for 10 min and washed in PGM. To stain callose, 0.1% aniline blue was added to cultured pollen tubes. To evaluate the distribution of demethylesterified pectin, pollen tubes were stained with 0.01% (w/v) ruthenium red for 5 min and washed in PGM ^5^. To assess cell membrane integrity, 2.5 μg/ml CellMask ™ Plasma Membrane Stain (Invitrogen) was applied for 5 min followed by three washes in PGM.

Pectin localization in the pollen tube cell wall was examined using JIM5, JIM7, and LM16 antibodies ^51^. Immunolabeling was conducted after fixing pollen tubes in freshly prepared 1.5% paraformaldehyde in PBS (pH 6.9) for 1.5 h at room temperature, followed by three rinses in PBS. Pollen tubes were treated with a 1:10 dilution (in PBS) of antibodies at room temperature for 2 h, followed by three washes in PBS. Subsequently, they were incubated with a 1:100 dilution of Alexa488-conjugated secondary antibodies at 4°C overnight and washed three times in PBS.

### Confocal microscopy and data analysis

Microscopy was conducted using the Leica STELLARIS 8. The following filters were used: for FM4–64, 543 nm excitation and a 560 nm long-pass emission filter; for Alexa 488, 488 nm excitation and a 515–565 nm emission filter; for aniline blue, 405 nm excitation and a 450– 490 nm bandpass emission filter; and for CellMask™ Plasma Membrane Stain, 561 nm excitation and a 586–614 nm bandpass filter. Confocal images were analyzed using Fiji (v. 2.14.0) and processed in Microsoft PowerPoint. A minimum of 100 independent pollen tubes were evaluated for each experiment to establish consensus phenotypes. Data are presented as means ± standard deviations (SDs). Pairwise comparisons were conducted using Student’s *t*-test to identify significant between-group differences. Statistical analysis was performed using Microsoft Excel software (v. 2017)

## Data Availability

The sequences of the genes used in this study were listed in Table S2 and can be accessed in the *Torenia* cDNA database (http://dandelion.liveholonics.com/torenia/).

## Acknowledgments

The authors gratefully acknowledge financial support from the MEXT Scholarship with Embassy Recommendation via the China Scholarship Council to X.J. This work was also supported by grants from the Japan Society for the Promotion of Science (21K15119 to A.G.M., 22H05172, 22H05178 to S.O., and 21K18235, 22H04980, 22K21352 to T.H.) and the Japan Science and Technology Agency (JPMJCR20E5 to T.H.).

## Authors’ Contributions

X.J., A.G.M., S.O., and T.H. designed the project. X.J. and A.G.M. performed most of the cytology experiments. X.J. conducted the bioinformatics analysis. X.J., S.O., and T.H. analyzed the data. X.J. and T.H. drafted the manuscript.

