## Supplementary Figures and Movies for "Vesicle-Mediated Regulation of Pollen tube Growth in *Torenia fournieri*"

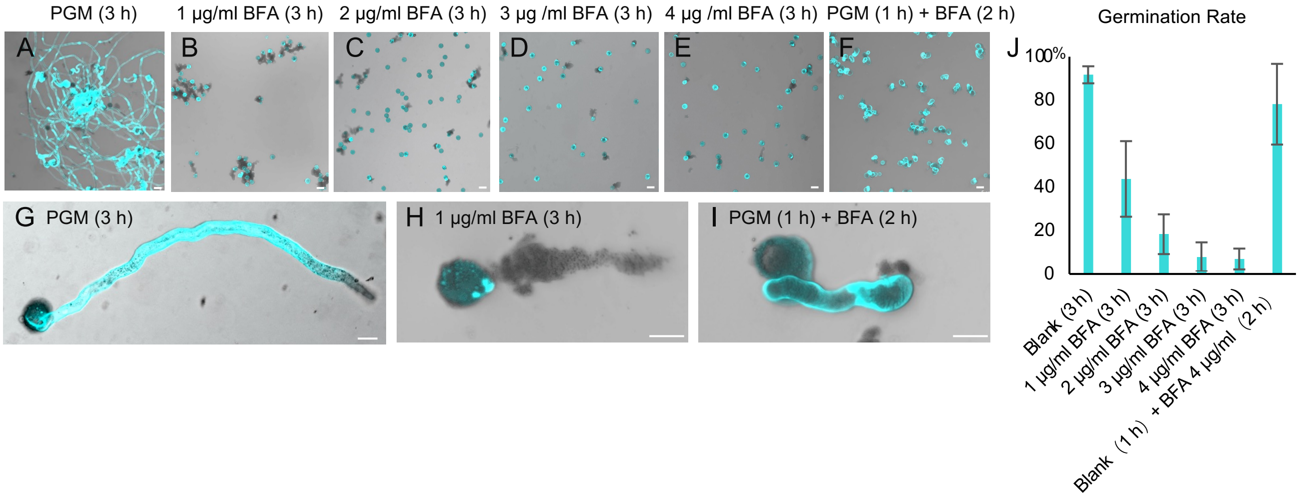


**Fig. S1. Effects of BFA at the indicated concentrations on pollen germination and pollen tube growth**

**A-F** *T. fournieri* pollen tubes cultured in PGM for 3 h (**A**); PGM containing 1 μg/ml BFA (**B**), 2 μg/ml BFA (**C**), 3 μg/ml BFA (**D**), and 4 μg/ml BFA (**E**) for 3 h; and Blank for 1 h followed by 4 μg/ml BFA for 2 h (**F**). **G-I** Morphology of pollen tubes cultured in PGM for 3 h (**G**), 1 μg/ml BFA for 3 h (**H**), and in Blank for 1 h followed by 4 μg/ml BFA for 2 h (**I**). **J** Pollen tube germination rate according to BFA concentration. Scale bars: 10 μm.

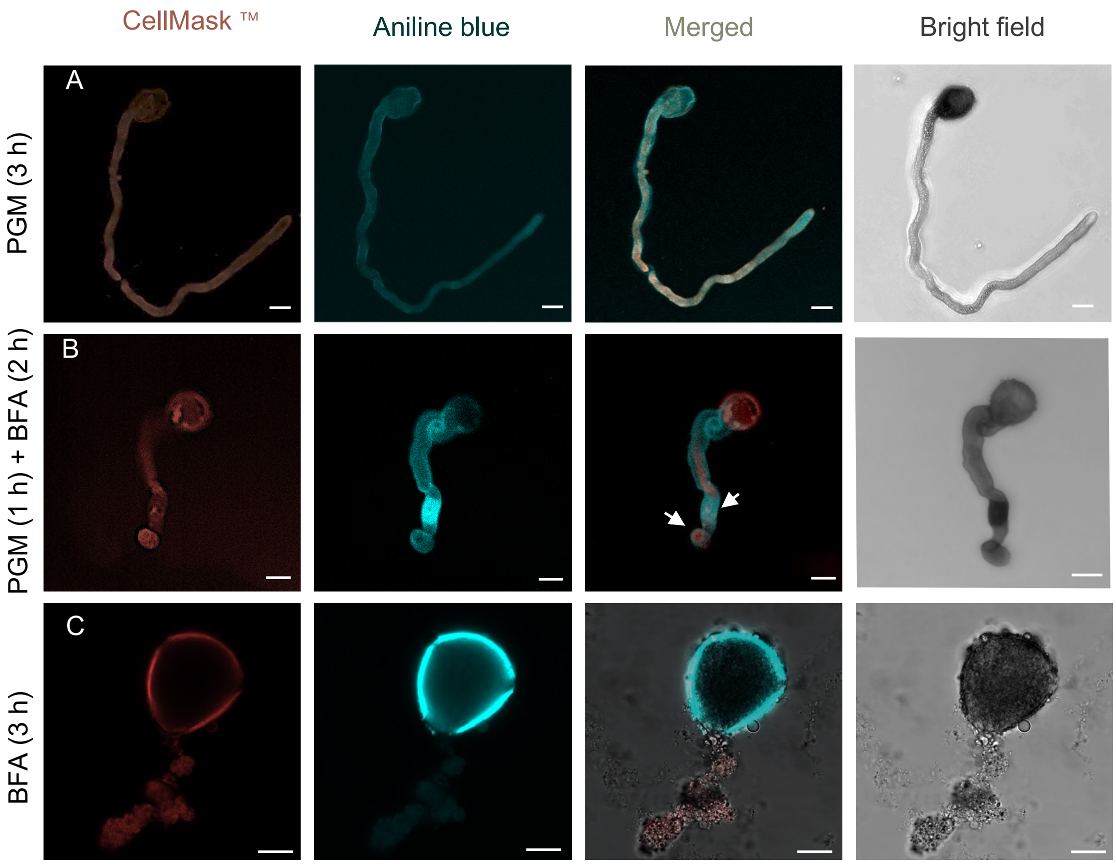


**Fig. S2. Cytochemical labeling of cell membrane and cell wall components**

**A** Images of a pollen tube cultured in PGM (3 h) stained with CellMask™ and aniline blue; a merged image and bright-field image are also shown. Bright-field image of the morphology of the control pollen tube. **B** Images of pollen tubes cultivated in PGM (1 h) and BFA (2 h) stained with CellMask™ and aniline blue; a merged image and bright-field image are also shown. Arrowheads indicate the abnormal callose distribution after PGM (1 h) and BFA (2 h) treatment. **C** Images of pollen tubes cultivated in BFA (3 h) stained with CellMask™ and aniline blue; a merged image and bright-field image are also shown. Scale bars: 10 μm.

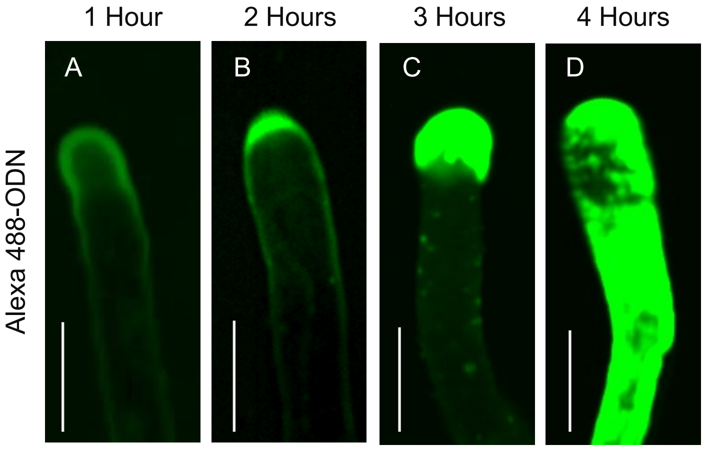


**Fig. S3. Uptake of antisense ODNs into pollen tubes cultivated in PGM for 4 h**

The distribution of Alexa 488-labeled ODN in pollen tubes after 1 (**A**), 2 (**B**), 3 (**C**), and 4 (**D**) hours of incubation. Scale bars: 10 μm.

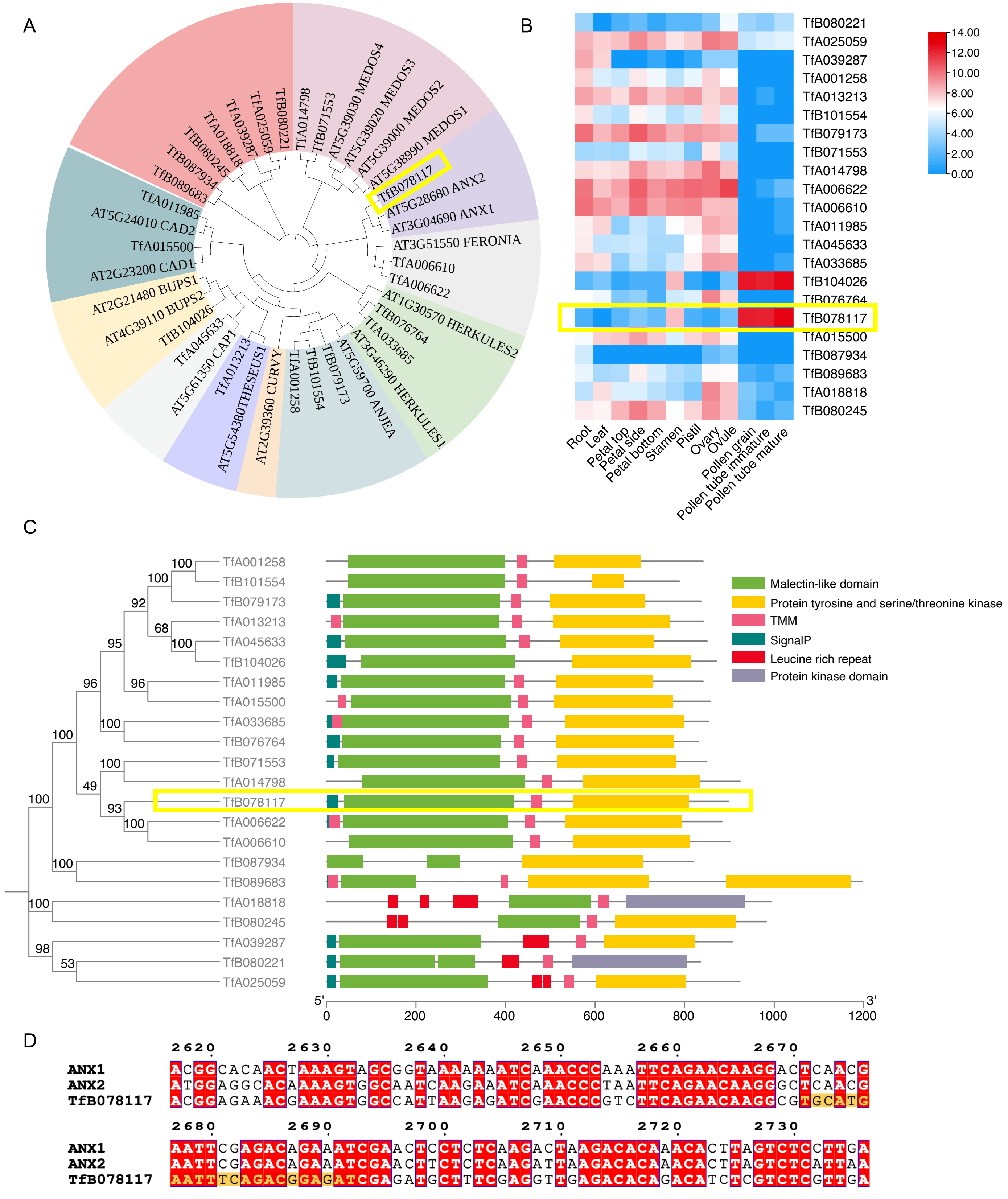


**Fig. S4. Genome-wide investigation of the *CrRLK1L* gene family and its expression profile in *T. fournieri***

**A** Evolutionary relationships of CrRLK1Ls in *A. thaliana* and *T. fournieri*. Seventeen *A. thaliana* CrRLK1L protein sequences were obtained from the TAIR website and 22 CrRLK1L proteins were identified in *T. fournieri*; these had malectin (pfam: TfB080671) and kinase (pfam: PF07714) domains. **B** Expression profile of *TfCrRLK1L* family members. **C** Domain structure of *TfCrRLK1L* family members. TfB078117, which had the closest evolutionary relationship to *ANX1* and *ANX2*, is highly expressed in pollen tubes and has a complete signal peptide and transmembrane region. Consequently, it is considered a homolog of *ANX* in *T. fournieri* and designated as *TfANX*. Yellow square, *TfB078117*. **D** Alignments of *TfB078117* with *FERONIA*, *ANX1* and *ANX2*, the ODN design site is highlighted in yellow.

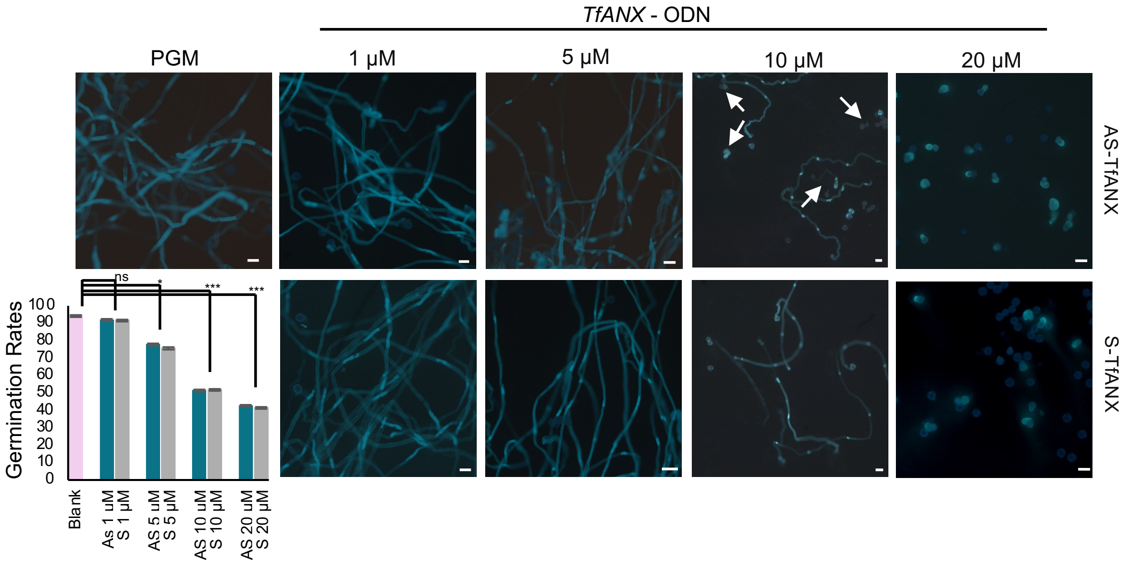


**Fig. S5. Concentration-dependent effects of AS-ODNs to *TfANX* on pollen germination**

Germination of pollen grains incubated for 3 h was examined in PGM containing the indicated concentrations of AS- and S-ODN. Concentration-dependent effects of ODNs on the pollen germination rate (n > 200). Arrows indicate abnormally developed pollen tubes. * P < 0.05, *** P < 0.001 from the control. Scale bars: 10 µm. Data are from ≥ 200 pollen tubes.

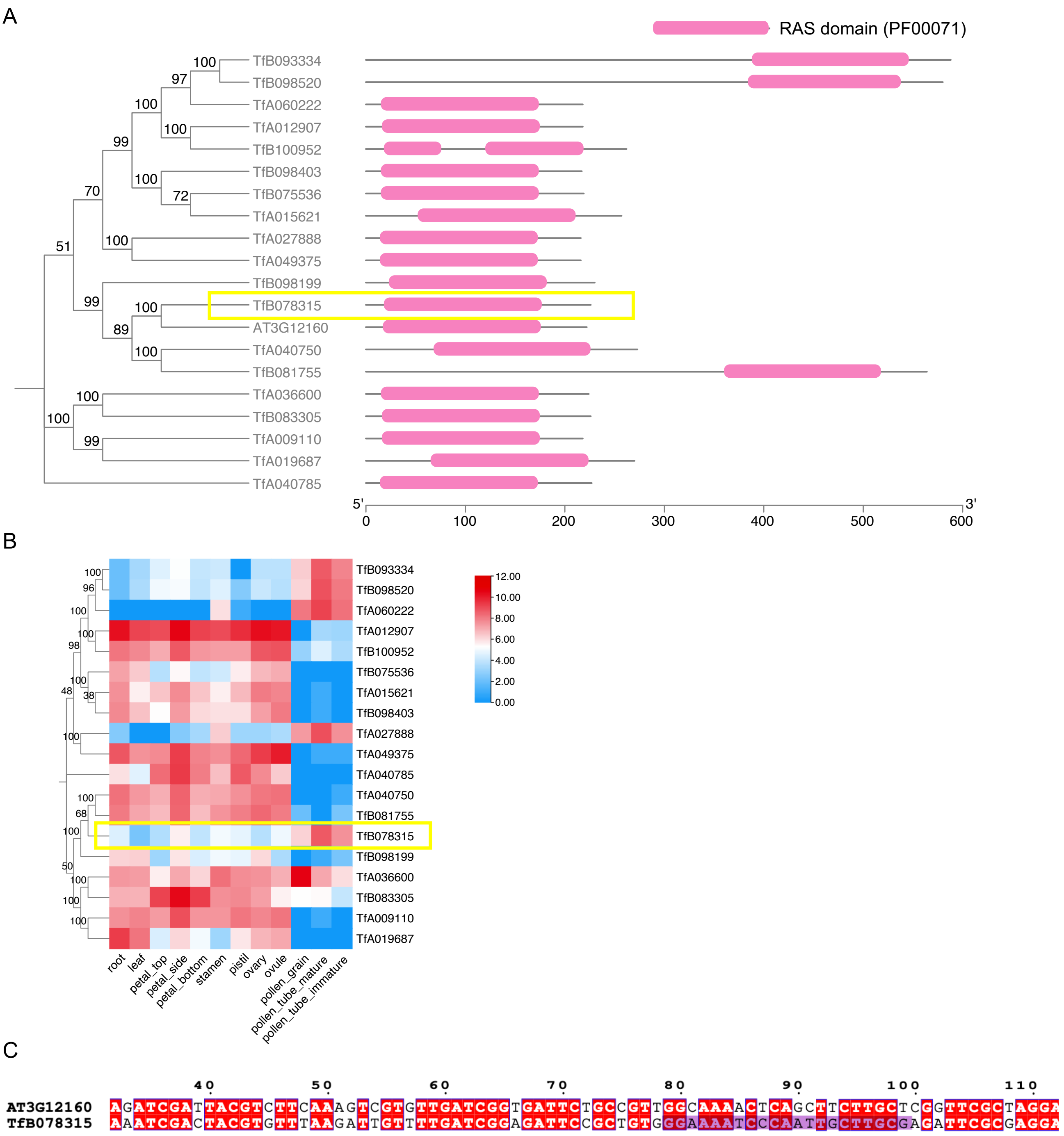


**Fig. S6. Genome-wide investigation of *RABA4D* homologs and their expression profiles in *T. fournieri***

**A** Evolutionary relationship of RABA4D and their domain structure in *A. thaliana* and *T. fournieri*. *A. thaliana* RABA4D protein sequences were obtained from the TAIR website; 19 RABA4D homologs had an RAS domain (pfam: PF00071). **B** Expression profile of *RABA4D* homologs. **C** Alignment of *TfB078315* with *RABA4D*; purple, ODN site. *TfB078315*, which had the closest evolutionary relationship to *RABA4D*, is highly expressed in mature pollen. Consequently, it was considered a homolog of *RABA4D* in *T. fournieri* and was designated *TfRABA4D*. Yellow square indicates the location of TfB078315.

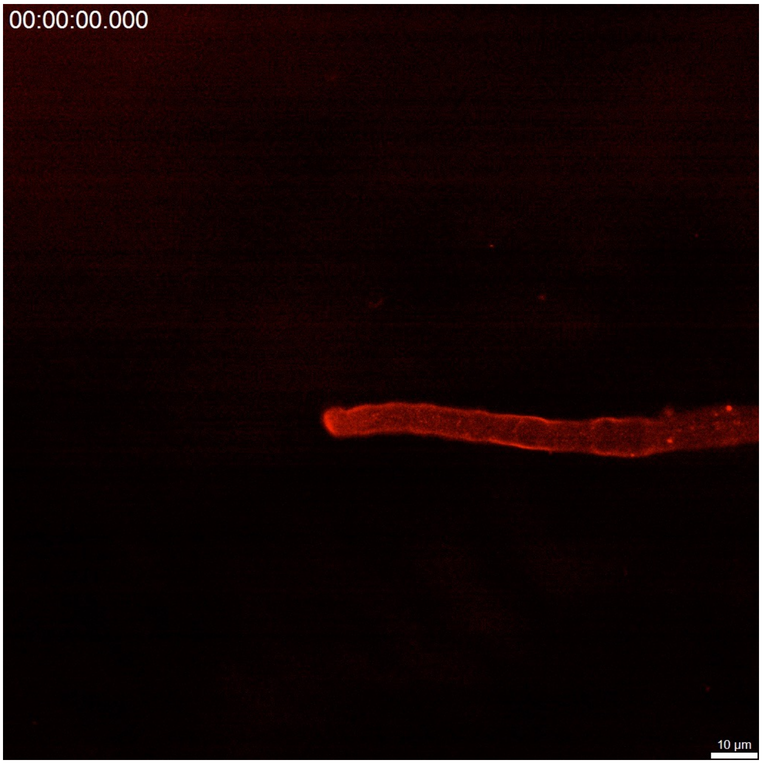


**Movie S1. Time-lapse recording of FM4-64 uptake in pollen tubes cultured in PGM (3 h)**

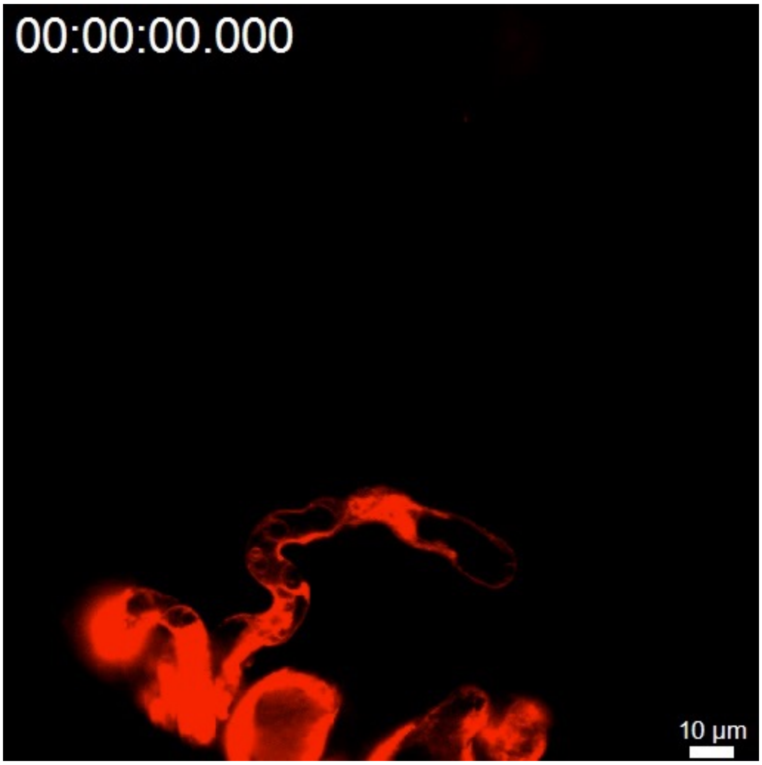


**Movie S2. Time-lapse recording of FM4-64 uptake in pollen tubes cultured in PGM (1 h) and BFA (2 h)**
